# Transcriptional differences between repeatedly recultured and freshly induced embryogenic calli reveal candidate developmental regulators

**DOI:** 10.64898/2026.10.25.761732

**Authors:** Hrishikesh Ingole, Gautam Saripalli, Sachin Rustgi

**Author notes:** Corresponding author: Dr. Sachin Rustgi.

## Abstract

Peanut (*Arachis hypogaea* L.) is an economically important oilseed and food legume, yet genetic transformation remains challenging because of genotype dependence and the lengthy tissue-culture and regeneration process. A better understanding of the molecular mechanisms underlying callus development and proliferation could help identify candidate morphogenic genes and developmental regulators (DRs) for peanut improvement. Here, we used RNA sequencing to compare embryogenic calli from the cultivar Georgia Green that had been maintained for three months (new callus; NC) or repeatedly subcultured for four years (old callus; OC). The two callus types displayed distinct morphologies: NC was pale cream, expanded, and cauliflower-like, whereas OC was more yellow, compact, and nodular. We identified 1,141 genes significantly upregulated in NC and 2,086 in OC (adjusted *p* < 0.05; log fold change > 2). Transcription factors were prominent among 13 categories of differentially expressed genes, including members of the AP2/ERF, WOX, GRAS, LBD, and NAC families, several of which are known DRs in other crops. qRT-PCR analysis of seven selected genes strongly corroborated the RNA-seq results (*r* = 0.944; *R*^2^ = 0.891). Highly upregulated candidates included C2H2/BED-type zinc-finger proteins, calreticulin/calnexin, a glycine-rich protein, a late embryogenesis abundant protein, glycoside hydrolases, and SWEET sugar transporters. These findings provide a resource for investigating callus development in peanut and prioritize candidate genes for functional validation and future improvement of transformation and regeneration systems.

## Introduction

Peanut (*Arachis hypogaea* L.) is an important legume crop primarily used for oil production (Huang et al., 2022). Other uses include peanut butter, snacks such as roasted peanuts, and as an extender in meat-based products, soups, and desserts (Arya et al., 2015). Its segmental allotetraploid genome and prevalence of homoeologous recombination make peanut a complex crop species for genetic analysis (Kochert et al., 1996; Bertioli et al., 2019; Ingole et al., 2026). Nevertheless, recent years have seen substantial advances in understanding the genomics of this complex crop species and the availability of genomic and genetic resources (Liu et al., 2026; Yang et al., 2026; Zhao et al., 2025; Bertioli et al., 2019; Zhuang et al., 2019; Lu et al., 2018). These developments have enhanced our understanding of the genetic architecture of agronomic and developmental traits.

The first reference-level peanut genome assemblies were published in 2019 (Bertioli et al., 2019; Zhuang et al., 2019), followed more recently by pan-genomes (Liu et al., 2026; Wang et al., 2025; Zhao et al., 2025). These assemblies revealed that the cultivated peanut genome comprises chromosome sets derived from ancestral species that have undergone several evolutionary changes, including mobile-element activity, deletions, and homoeologous recombination. At the same time, efforts to improve developmental transcriptome datasets have yielded expression profiles across diverse peanut tissues and developmental stages, including flowers, pegs, pods, seeds, roots, nodules, and vegetative tissues. Together, genomic and transcriptomic resources have enabled more precise interpretation of RNA-seq results in peanut—not only by identifying differentially expressed genes, but also by placing these candidate genes in broader developmental and tissue-specific contexts (Clevenger et al., 2016; Bertioli et al., 2019). These resources have also enabled the development of genomic and genetic tools for studying important traits, including oil content (Xue et al., 2025b), allergens (Conner et al., 2024; Alam et al., 2026), and responses to biotic and abiotic stresses, such as late leaf spot and drought (Gangurde et al., 2024; Abady et al., 2024).

Understanding the genetic regulation of important traits is essential for improving peanut. Although conventional molecular markers are already in use (Varshney et al., 2014; Deshmukh et al., 2020), CRISPR-based gene editing, overexpression, and RNAi technologies have been used only to a limited extent in peanut. This is largely attributable to the lack of efficient regeneration protocols, which is primarily due to the crop’s tetraploid genome and recalcitrance to regeneration. Nevertheless, studies have evaluated different explants—including leaflets, cotyledons, immature cotyledons, embryonic axes, and hypocotyls—to establish successful regeneration protocols (see Phogat et al., 2024, for a review). A few studies have also reported Agrobacterium-mediated transformation using strains such as LBA4404, GV2260, EHA101, EHA105, GV3101, and C58 (Rohini and Rao, 2000; Anuradha et al., 2006; Kim et al., 2009), as well as *in planta* particle bombardment (Joshi et al., 2005; Chu et al., 2013). CRISPR-Cas9 genome editing in peanut has been reported for developing fragrant peanut (Xu et al., 2025), high-oleic-acid lines through editing of *FAD2* genes (Yuan et al., 2019), and lines with reduced allergen content (Conner et al., 2024; Alam and Rustgi, 2025). However, progress remains limited by low tissue-regeneration efficiency following callus induction, the small number of peanut cultivars to which these methods have been applied, and lengthy timelines that often exceed a year from seed to seed.

Callus regeneration is an integral part of *in vitro* plant tissue culture protocols. Recent studies have shown that callus regeneration is genotype-dependent, which can limit the development of reliable transformation systems across peanut germplasm. Genome-wide association studies (GWAS) have revealed genetic variation in callus induction among peanut accessions (Luo et al., 2024), while QTL mapping has identified loci affecting embryogenic callus formation and regeneration in cereals and legumes (Potokina and Sushchenko, 2025). These findings highlight the contribution of genotype-specific genetic factors to *in vitro* recalcitrance. Similarly, the effects of different plant growth regulator combinations on peanut regeneration have been demonstrated in the cultivar Georgia-12Y (Khatri and Joshi, 2024). That study highlighted the critical roles of plant growth regulators in callus formation, somatic embryogenesis, shoot organogenesis, and successful plant establishment. Traditional tissue culture systems rely heavily on combinations of auxins and cytokinins to induce callus formation from explants. Several studies have also reported that callus formation is regulated by hormone signaling, transcriptional control, cell-cycle activation, and epigenetic changes (Skoog and Miller, 1957; Fan et al., 2012; Ikeuchi et al., 2013; Mehraj et al., 2025).

Limitations in callus regeneration and plant tissue culture have prompted research into the use of developmental regulators (DRs) to enhance regeneration from callus tissue or induce plant organs *de novo* (Jiang et al., 2024). DRs were first identified in *Arabidopsis* in the 1990s (Laux et al., 1996; Lotan et al., 1998). The role of WUSCHEL in shoot and floral meristem development was first demonstrated by Laux et al. (1996). In 1998, another important DR, *leafy cotyledon 1* (LEC1), was identified and shown to be involved in embryo development and maturation (Lotan et al., 1998). In the 2000s, many other DRs were identified, primarily in *Arabidopsis*. These included transcription factors such as GRAS, LBD, PLETHORA, and AP2, with demonstrated roles in callus formation and regeneration, as well as root and shoot development. By the late 2000s, the use of DRs to support shoot, root, and callus regeneration had expanded to other dicots, including *Brassica*, soybean, and sunflower (Kong et al., 2020), and monocots, including maize, rice, and wheat (Lowe et al., 2016; Debernardi et al., 2020). This helped popularize DRs in modern plant tissue culture across many important plant species. Despite its importance as a food crop, peanut remains relatively underexplored in this area (Gu et al., 2026; Tang et al., 2024; Liu et al., 2021; Tang et al., 2018), highlighting the need to identify DRs that can accelerate callus regeneration and address key limitations. Studies in peanut have identified roles for LEC1 in seed development (Tang et al., 2018, 2024) and embryo and cotyledon development (Tang et al., 2024); *AhWOX13* and *AhWOX20* in adventitious shoot regeneration (Gu et al., 2026); and *AHL23* in leaf development (Liu et al., 2021). However, with the exception of *AtLEC1* (Tang et al., 2018), the remaining genes, though identified in peanut, have been functionally validated in *Arabidopsis*. This underscores the need to characterize them further in peanut.

The present study is an effort to understand the molecular mechanisms underlying embryogenic callus development in peanut and to identify putative DRs that may enhance regeneration after further validation. To this end, we performed RNA-seq analysis of long-term-maintained and newly induced peanut calli and identified differentially expressed genes (DEGs). Some of these may be candidate DRs because of their known roles in other crops or their significantly higher expression in embryogenic calli. A recent study also reported differential gene expression in peanut (Xue et al., 2025a). However, that study focused on identifying peanut-specific callus promoters that could restrict CRISPR component expression to callus tissue and prevent unintended expression in regenerated plants. We further validated the expression of several candidates identified in the present study using qRT-PCR. Overall, to our knowledge, this is the first preliminary study to report on the molecular mechanisms of embryogenic callus development in peanut and to identify putative DRs for callus induction and regeneration.

## Materials and Methods

### Plant material, callus induction and maintenance

‘Georgia Green’ (Reg. no. CV-55, PI 587093), a popular, tissue culture-friendly, small-seeded, runner-type peanut variety resistant to Tomato spotted wilt virus (TSWV), was used for callus induction in this study. Seeds were obtained from North Carolina State University (NCSU). For callus induction, seeds were surface-sterilized, and embryos were excised according to the procedure described by Chu et al. (2019). The excised embryos were kept in the dark on callus induction medium and sub-cultured every 15 days. The callus induction medium comprised 5.24 g L ¹ Murashige and Skoog (MS) salts (MS basal salts mixture; PhytoTechnology Laboratories, Lenexa, KS, USA), 15 g L ¹ sucrose, 2 g L ¹ Phytagel, 1 mL L ¹ picloram, and PPM at the manufacturer’s recommended concentration (Plant Cell Technology, Washington, DC, USA). After two months, the calli were transferred to maintenance medium consisting of callus induction medium supplemented with 0.3% glutamine (w/v). They were sub-cultured on the same medium every 21 days for four years (2021–2025). Callus subcultures were maintained at 28 °C under a 16 h light/8 h dark photoperiod, with 1,000 ppm CO, cool-white, fluorescent lighting at an irradiance of 60 μmol m ² s ¹, and 50% relative humidity.

### Total RNA extraction, RNA-seq and Data analysis

Two callus culture treatments were used for RNA-seq analysis: cultures maintained for four years, hereafter referred to as old callus cultures (OC), and cultures maintained for three months, hereafter referred to as new callus cultures (NC). Total RNA was extracted from three biological replicates of each treatment using the Spectrum™ Plant Total RNA Kit (Sigma-Aldrich, Inc., St. Louis, MO, USA). RNA concentration was measured using a NanoDrop One spectrophotometer (Thermo Fisher Scientific Pvt. Ltd., USA), and RNA integrity was assessed by electrophoresis on a 1% TAE agarose gel. The six RNA samples (three biological replicates each for OC and NC) were sent to Azenta Life Sciences (USA) for library preparation and RNA sequencing.

### Library Preparation with PolyA selection and Illumina Sequencing

RNA was quantified using Qubit 3.0 Fluorometer (Life Technologies, Carlsbad, CA, USA) and RNA integrity was reassessed using Agilent TapeStation 4200 (Agilent Technologies, Palo Alto, CA, USA).

RNA-sequencing libraries were prepared using the NEBNext Ultra II RNA Library Prep Kit for Illumina according to the manufacturer’s instructions (NEB, Ipswich, MA, USA). Briefly, mRNA was enriched using oligo(dT) beads and fragmented for 15 min at 94 °C. First- and second-strand cDNA were then synthesized. The cDNA fragments were end-repaired and adenylated at their 3′ ends, and universal adapters were ligated to the fragments. Indexes were added, and the libraries were enriched by limited-cycle PCR. The sequencing libraries were validated using an Agilent TapeStation (Agilent Technologies, Palo Alto, CA, USA) and quantified using a Qubit 3.0 Fluorometer (Invitrogen, Carlsbad, CA, USA) and quantitative PCR (KAPA Biosystems, Wilmington, MA, USA).

The sequencing libraries were multiplexed and clustered on a flow cell using an Illumina NovaSeq instrument, according to the manufacturer’s instructions. Samples were sequenced in a paired-end 2 × 150 bp configuration. Image analysis and base calling were performed using NovaSeq Control Software (NCS). Raw sequence data (.bcl files) generated by the instrument were converted to FASTQ files and demultiplexed using Illumina bcl2fastq v2.20 software, allowing one mismatch for index sequence identification. Raw RNA-seq reads for all six samples were provided by Azenta Life Sciences and used for downstream analyses.

### Bioinformatics

The raw reads obtained from Azenta Life Sciences were trimmed with Cutadapt v5.2 to remove adapter sequences and low-quality reads. The cleaned reads were mapped to the peanut genome (arahy.Tifrunner.gnm2; Bertioli et al., 2019) using HISAT2 v2.0 with default parameters. The resulting SAM files were converted to BAM format, and gene-level read counts were quantified with featureCounts using the annotation corresponding to the *Arachis hypogaea* cv. Tifrunner reference genome. The unnormalized count matrix was analyzed in R using DESeq2 (v1.48.0). Counts were normalized using DESeq2-estimated size factors, and differential expression was assessed with a negative binomial generalized linear model. Genes with an adjusted P value < 0.05 and an absolute log fold change > 2 were considered significantly differentially expressed. Throughout the manuscript, genes with positive log fold-change values (NC relative to OC) were considered upregulated in NC, whereas genes with negative values were considered upregulated in OC.

### RGB-based analysis of callus pigmentation

Both OC and NC samples were photographed using a Canon DSLR camera (EOS Rebel T6), and the resulting images were analyzed using Batch Load Image Processor 1.1 (BLIP; Clemson University, Clemson, SC, USA). The image-processing workflow, including the software used to extract RGB values, was adapted from Renfroe-Becton et al. (2022). Red (R), green (G), and blue (B) values were compared between the NC and OC groups, with three samples per group. Each color was analyzed separately using a two-sided Welch’s independent-samples t-test, which allows for unequal variances between groups. P-values were adjusted across the three color comparisons using the Holm method, with an adjusted significance threshold of 0.05. The figure was generated in R and displays individual sample values alongside the mean ± standard deviation (SD). The same vertical-axis scale was used across all three panels to facilitate comparison.

### qRT-PCR analysis of selected genes

Expression of seven representative DEGs identified by RNA-seq analysis was further validated by qRT-PCR. The selected genes included three predicted to be highly upregulated in OC and four predicted to be highly upregulated in NC (Table 1).

Total RNA was extracted from cotyledonary leaves using the Sigma Total Plant RNA Kit. First-strand cDNA was synthesized from 1 µg of total RNA in a 20 µL reaction using the Thermo Scientific Maxima First Strand cDNA Synthesis Kit. RT-qPCR was performed using Bio-Rad SYBR® Green Master Mix on a Bio-Rad 5 Real-Time PCR System. *Ahadh3* was used as the reference gene (Xue et al., 2025a). Each sample was analyzed using three biological replicates and three technical replicates. Relative gene expression levels were calculated using the 2^−ΔCt^ method (Livak et al., 2002). Primer sequences used for RT-qPCR are listed in Table 1.

Concordance between RNA-seq and qRT-PCR results was assessed using log fold-change values for seven selected genes, expressed as NC relative to OC. RNA-seq and qRT-PCR values were plotted on the x- and y-axes, respectively, using the ggplot2 package in R, and gene labels were positioned using ggrepel. Pearson’s correlation coefficient (r) and its two-sided P value were calculated to evaluate the association between the two methods. A simple linear regression was fitted, and the coefficient of determination (R²) was reported. The regression line was displayed with its 95% confidence interval, and a dashed 1:1 reference line indicated equal fold-change estimates between methods.

### Chromosomal distribution of DEGs and physical mapping of selected candidate genes

The genomic distribution of the 3,227 DEGs listed in Table S4 was examined using gene coordinates from the *Arachis hypogaea* cv. Tifrunner reference annotation (gnm2.ann2). Chromosome lengths were obtained from the corresponding reference genome FASTA index. Chromosomes Chr01–Chr10 and Chr11–Chr20 were assigned to the A and B subgenomes, respectively. Each gene was positioned using the midpoint of its annotated start and end coordinates. Of the 3,227 DEGs, 3,221 mapped to the 20 chromosomes; the remaining six mapped to unplaced scaffolds and were excluded from chromosome-based plots.

Gene identifiers were extracted from Table S4 using the Python package openpyxl and matched to genomic coordinates in the Tifrunner gnm2.ann2 GFF3 annotation. Genes were assigned to non-overlapping 5-Mb chromosomal windows according to their genomic midpoints, and DEG counts were calculated using NumPy. Counts in shorter terminal windows were normalized to a 5-Mb equivalent. Chromosomes were arranged sequentially around a circular layout, with segment lengths proportional to chromosome lengths. DEG frequencies were represented by a yellow-to-red color gradient (Figure S1).

## Results

### Morphological changes during callus development

NC and OC showed distinct morphological characteristics. NC was pale cream in color and had a relatively expanded, irregular, cauliflower-like appearance. In comparison, OC was more yellowish and had a compact, nodular morphology with distinct, rounded structures. Thus, prolonged callus maintenance was associated with visible changes in pigmentation and tissue organization (Fig. 1).

**Figure 1.**
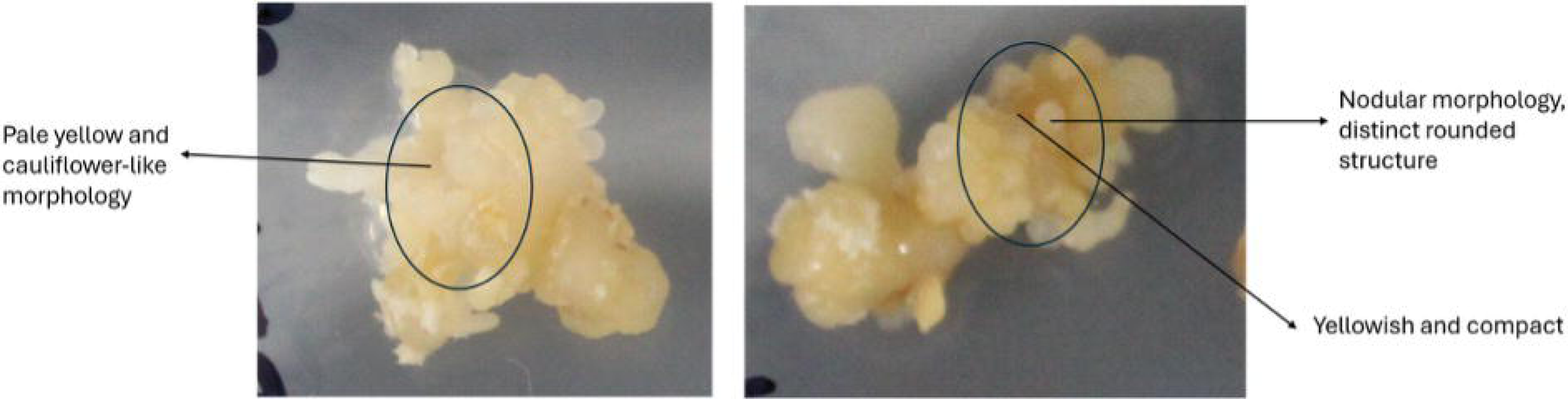
Representative morphology of newly developed callus (NC; left) and old callus (OC; right).

Comparisons of R, G, and B values between NC and OC revealed higher mean values for all three color channels in NC: 207.60 ± 3.50 for R, 185.67 ± 1.31 for G, and 157.36 ± 9.92 for B. The corresponding values for OC were 182.46 ± 7.65 for R, 157.36 ± 9.92 for G, and 111.49 ± 9.55 for B. The mean differences were 25.14, 28.31, and 29.33 intensity units for R, G, and B, respectively (Table S1).

All three comparisons narrowly met the significance threshold after Holm adjustment (adjusted p = 0.0497 for each color channel). Given the small sample size, these findings provide preliminary evidence of differences in RGB intensity between new and old calli.

### RNA-seq data, differentially expressed genes, and their chromosomal distribution

A total of 198,432,960 reads were obtained across the six samples. NC-Rep3 had the highest number of reads, while OC-Rep2 had the lowest. The mean number of reads per sample wa 33,072,160. Alignment to the peanut genome ranged from 78.35% in OC-Rep2 to 96.89% in OC-Rep1, with a mean alignment rate of 93.23% across all six samples (Table S2). All sample except OC-Rep2 showed alignment rates above 95%.

Analysis of the aligned reads identified 47,611 DEGs, of which 3,227 were significantly differentially expressed (adjusted p < 0.05 and absolute log fold change > 2; Table S4). Among these significant DEGs, 1,141 were upregulated in NC and 2,086 in OC (Table S3). Their distribution is shown in the volcano and MA plots (Figures 2a and 2b).

Chromosomal distribution of the 3,227 DEGs revealed their presence across all 20 peanut chromosomes, spanning both the A and B subgenomes. The observed gene density in 5-Mb windows was similar to the expected gene density based on known annotations (Tifrunner reference annotation; gnm2.ann2), although DEG frequency varied among windows (Figure S1). Twenty-seven candidates from the top 20 lists—eight upregulated in NC and 19 in OC—mapped to 13 different chromosomes.

**Figure 2.**
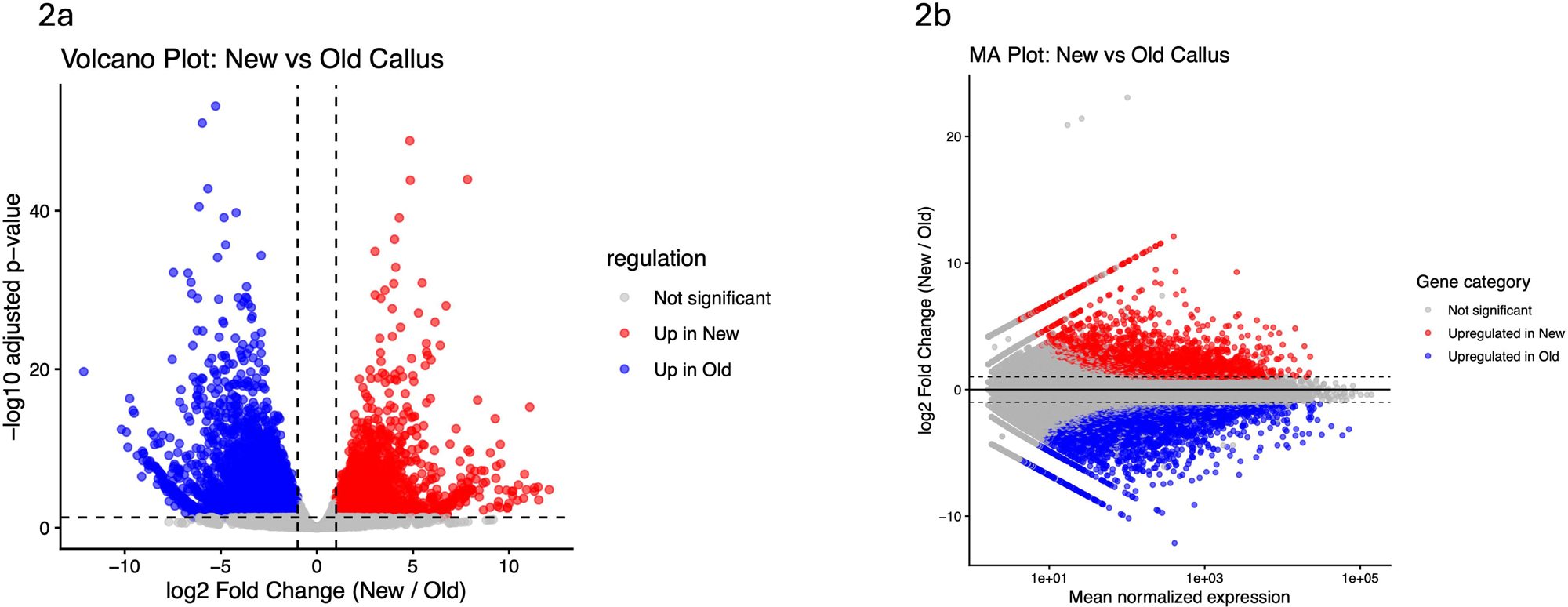
Transcriptomic comparison of new and old callus tissues. (a) Volcano plot showing differential gene expression between new and old callus. The x-axis shows log fold change (new/old), and the y-axis shows −log adjusted p-value. Red points indicate genes significantly upregulated in new callus, blue points indicate genes significantly upregulated in old callus, and gray points indicate non-significant genes. Dashed lines mark the significance thresholds of |log fold change| ≥ 1 and adjusted p < 0.05. (b) MA plot showing log fold change (new/old) versus mean normalized gene expression. Red, blue, and gray points indicate genes upregulated in new callus, genes upregulated in old callus, and non-significant genes, respectively.

### Functional annotation of DEGs

In total, the DEGs were classified into 13 categories based on available annotations (Table 2, Figure 3, and Table S4): (i) transcription factors, (ii) genes involved in metabolism, (iii) gene involved in the oxidative stress response, (iv) kinases, (v) genes involved in cell wall metabolism, (vi) transporters, (vii) disease resistance genes, (viii) important enzymes, (ix) seed storage proteins, (x) methyltransferases, (xi) growth response genes, (xii) photosynthesis-related genes, and (xiii) genes in the miscellaneous category. Genes in the miscellaneous category were the most abundant (791; 24.5%), followed by transcription factors (463; 14.3%), genes involved in metabolism (329; 10.2%), and genes involved in the oxidative stress response (302; 9.4%). Important genes identified in each category are listed in Table S4, and the 20 DEGs with the most extreme expression values are listed in Table 3.

**Figure 3.**
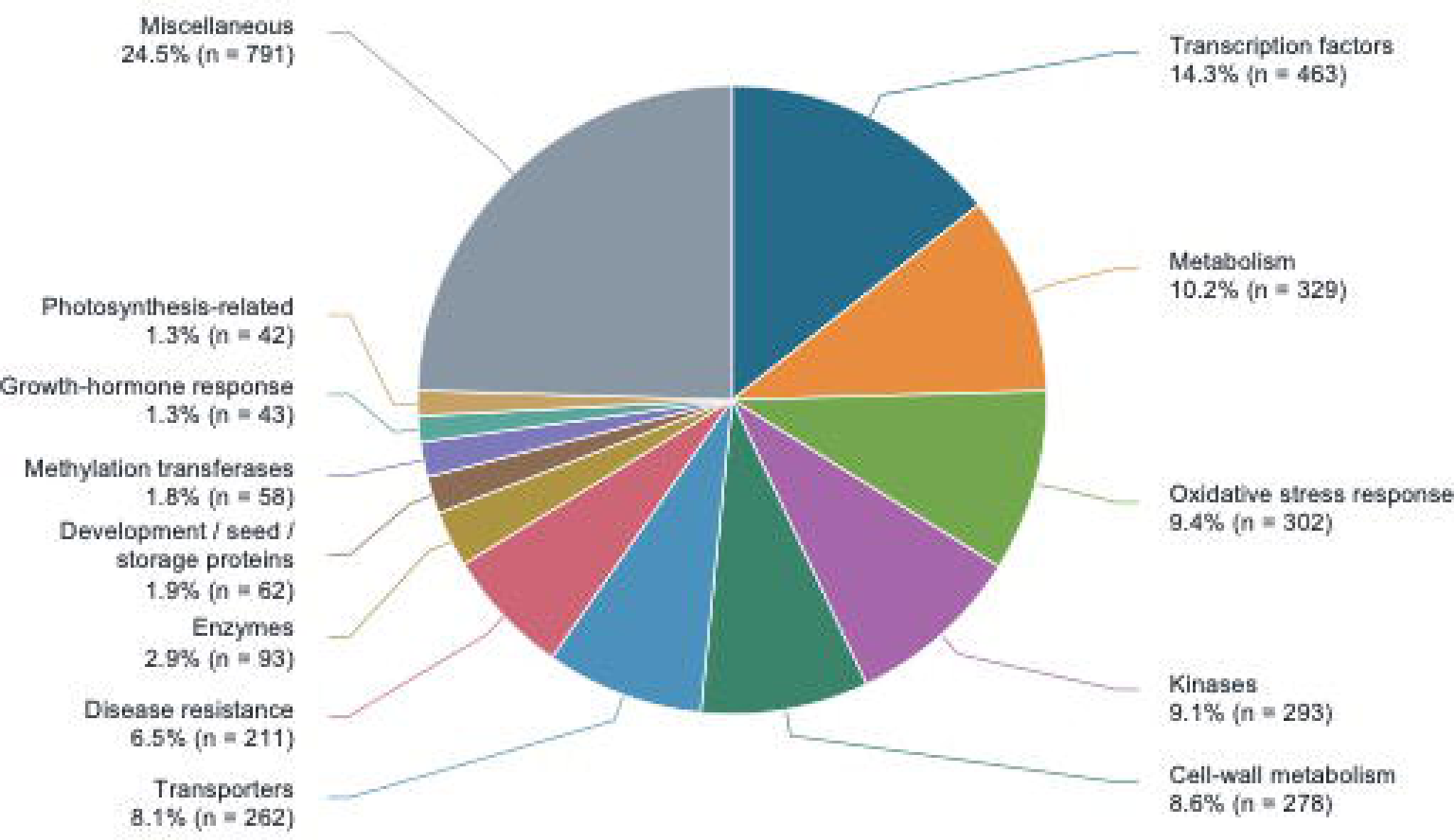
Functional classification of differentially expressed genes in NC versus OC. The size of each pie-chart sector represents the proportion of genes assigned to that category, and the value within each sector indicates the number of genes. The miscellaneous category includes genes that could not be assigned to any of the predefined functional groups.

### Gene ontology and enrichment

Gene ontology (GO) analysis classified the annotated DEGs into three major categories: biological process, molecular function, and cellular component (Figure 4). Predominant biological process terms included regulation of biological processes (GO:0050789), biological regulation (GO:0065007), localization (GO:0051179), and signaling (GO:0023052). Major molecular function terms included catalytic activity, binding, transporter activity, transcription regulator activity, oxidoreductase activity, protein binding, and structural molecule activity. Prominent cellular component terms included cell, cell part, organelle, plastid, nucleus, cytoplasm, extracellular region, and membrane-associated components.

GO enrichment analysis identified functions related to stress responses, carbohydrate and cell wall processes, photosynthesis, and oxidative stress responses as significantly enriched (P < 0.05; Figure 5). Enrichment of terms related to responses to oxidative stress and biotic stimuli suggests differences in stress-associated transcription between newly induced and long-term-maintained callus. These differences may reflect responses to callus induction, repeated subculturing, or prolonged exposure to *in vitro* culture conditions. However, because the enrichment analysis used the combined DEG set, these processes cannot be attributed specifically to either callus stage.

Enrichment of terms related to cell wall modification, cell wall organization, and xyloglucan metabolism indicates transcriptional differences associated with cell wall remodeling. Together with the carbohydrate-related processes, these findings suggest changes in carbon utilization and cell wall dynamics that may contribute to differences in callus growth, texture, and friability. Establishing these relationships would require direct physiological and cell wall measurements.

Photosynthesis-related GO terms also suggest differences in plastid-associated functions and metabolic or differentiation states between the two callus types. Overall, GO classification and enrichment analyses identified coordinated differences in regulatory, metabolic, stress-response, and structural processes associated with the two callus stages.

**Figure 4.**
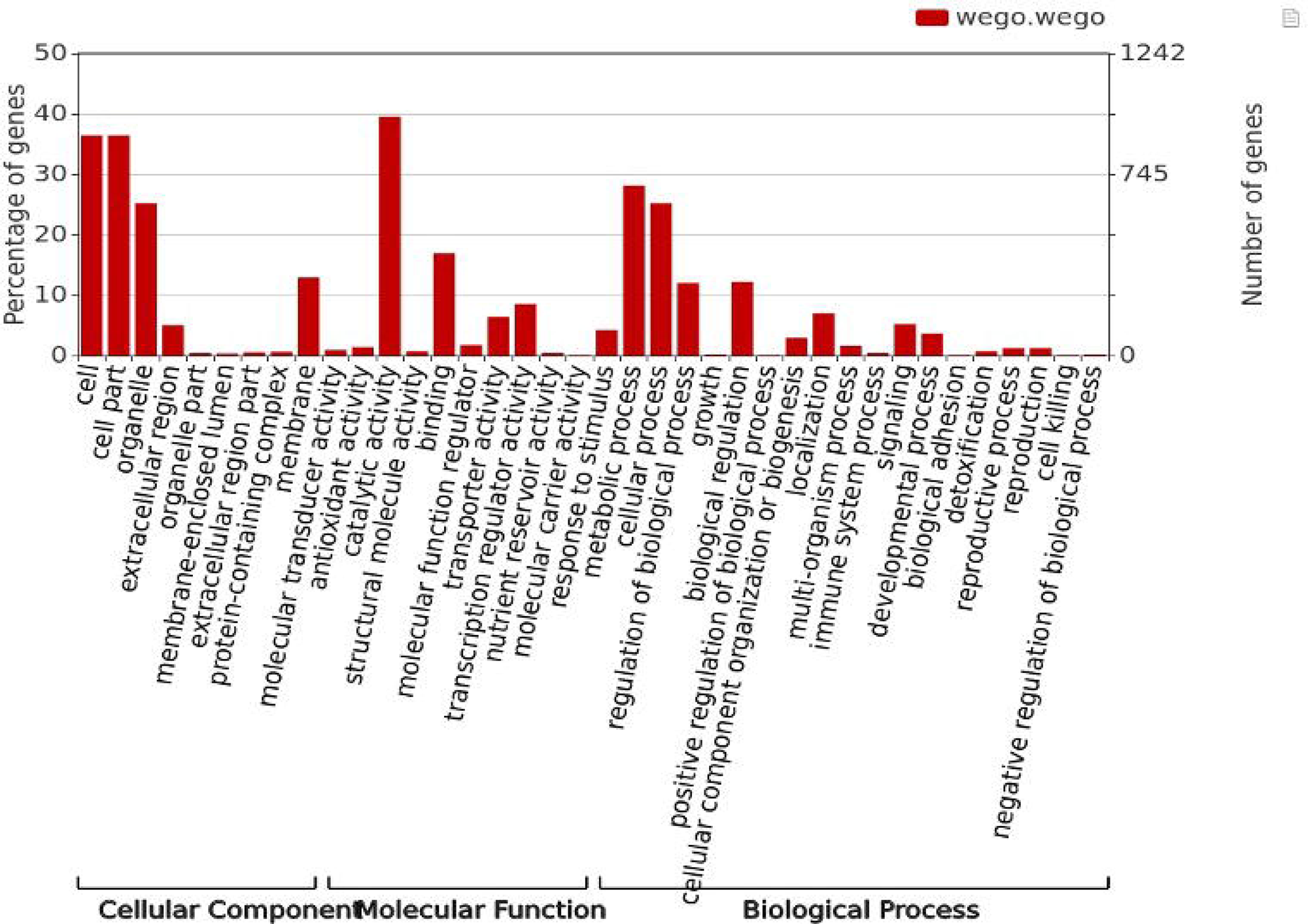
Gene Ontology classification of differentially expressed genes in newly induced versus long-term-maintained peanut callus.

**Figure 5.**
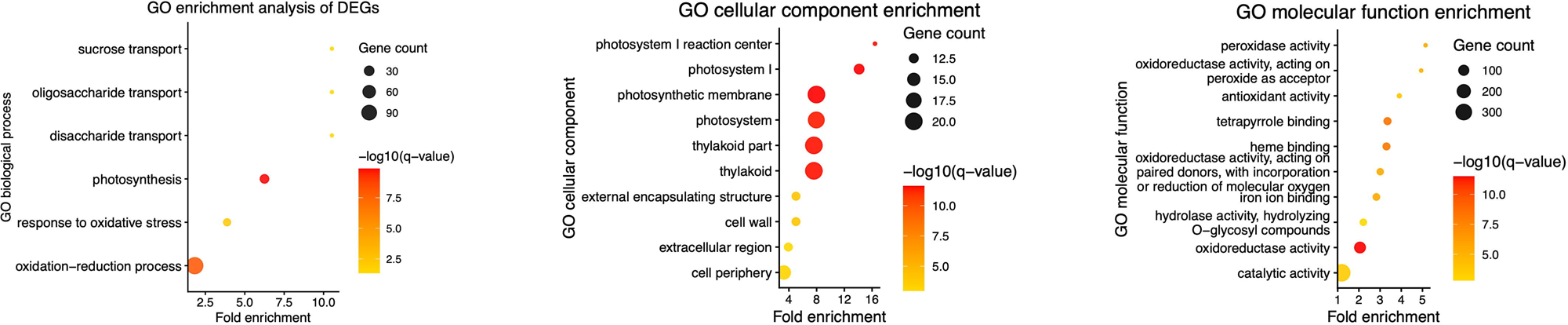
Enriched Gene Ontology terms among differentially expressed genes, grouped by biological process, cellular component, and molecular function.

### Validation of selected gene expression profiles by qRT-PCR

qRT-PCR analysis of seven selected DEGs in NC and OC revealed significant differences in expression between the two treatments. Relative fold-change values ranged from −5.80 for Ah13g128600 to 10.82 for Ah06g297300 in NC relative to OC. Four genes (Ah03g400100, Ah06g297300, Ah17g087800, and Ah02g299800) showed higher expression in NC, whereas the other three (Ah13g128600, Ah13g032500, and Ah20g092000) showed higher expression in OC (Table S5; Figure 6).

Comparison of the qRT-PCR and RNA-seq fold-change values revealed a strong positive correlation (Pearson’s r = 0.944; R² = 0.891), although the magnitude of the fold changes differed between the two methods (Table S5; Figure 6). Overall, these results support the consistency of the expression directions identified by RNA-seq.

**Figure 6.**
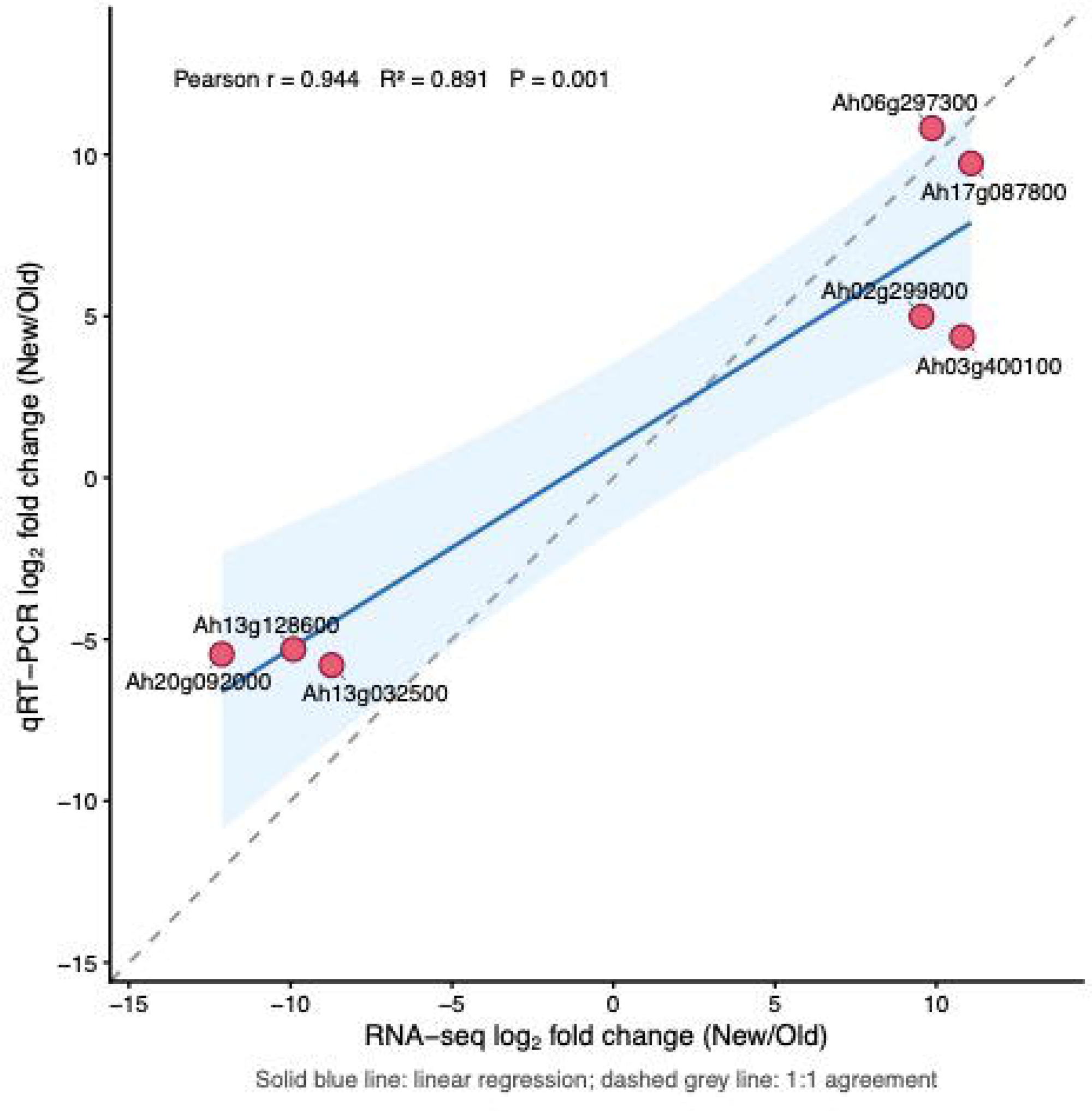
Comparison of expression changes measured by RNA-seq and qRT-PCR for seven selected genes in peanut callus.

## Discussion

*In vitro* plant regeneration is a complex process involving a series of biochemical, morphological, and molecular changes. Tissue culture generally involves somatic embryogenesis or direct regeneration. In polyploid crops such as peanut, regeneration through somatic embryogenesis is challenging for several reasons (Chengalrayan et al., 1997, 1998; Khatri and Joshee, 2024). Growth regulators, particularly auxins and cytokinins, are conventionally used in different combinations and proportions to promote regeneration in crops, including peanut.

DRs have also been widely studied for this purpose, primarily in *Arabidopsis* and a few other crops. However, their application in peanut has rarely been demonstrated, with only a few exceptions (Tang et al., 2018, 2024; Liu et al., 2021; Gu et al., 2026). Although these studies report the use of DRs in peanut, most of their findings have been validated in *Arabidopsis*. One exception is Tang et al. (2018), who studied *AtLec1*, a gene identified in *Arabidopsis* and found that it is involved in peanut seed development. The present study aims to identify potential DRs in peanut that may enhance the capacity of callus to regenerate shoots and/or roots. Identifying DRs requires an understanding of the molecular mechanisms underlying callus development. Therefore, we performed RNA-seq analysis on callus maintained *in vitro* and continuously subcultured for four years, as well as on newly induced callus. We identified DEGs between the two callus types and discuss their implications for developmental regulation.

### Distinct morphological changes in old and new calli

The two types of callus exhibited the expected morphological differences. Newly induced callus (NC) was pale cream in color, relatively expanded, and irregular in appearance, resembling a cauliflower. In contrast, long-term-maintained callus (OC) was more yellowish and had a compact, nodular morphology with distinct, rounded structures. These observations suggest that prolonged callus maintenance was associated with visible changes in pigmentation and tissue organization (Fig. 1). Similar observations have been reported in *Arabidopsis* and Citrus, where embryogenic callus was white and rich in organelles, whereas aged callus was pale brown and contained large vacuoles (Wang et al., 2026; Lee et al., 2016).

In our study, both OC and NC were derived from mature seed embryos and were embryogenic. Prolonged *in vitro* callus maintenance can be accompanied by epigenetic reprogramming and genetic alterations that contribute to somaclonal variation (Ma et al., 2015; Li et al., 2019; Wang et al., 2022; Smith and Street, 1974). Epigenetic modifications may improve regenerability in some crops, such as cotton, or reduce callus regeneration capacity in others, such as carrot. In carrot, cytological changes have also been observed alongside epigenetic and somaclonal variation. In cotton, improved regeneration was primarily attributed to a process known as Successive Regeneration Acclimation (SRA), which involved targeted epigenetic modifications, specifically the remodeling of DNA methylation and chromatin accessibility. This technique resulted in the development of the highly regenerative elite line Jin668 from the less-regenerable line Y668 (Li et al., 2019). Furthermore, repeatedly subcultured cells may also exhibit changes in ploidy levels (Smith and Street, 1974).

Although epigenetic variation is one of the major changes associated with repeated subculturing, the morphological differences between the two callus types may also be associated with the differential expression of numerous other genes. These include genes involved in cellular dedifferentiation and in the totipotency and regenerative capacity of somatic embryos to give rise to different tissues. This possibility is supported by earlier studies in *Arabidopsis* reporting large-scale upregulation of transcription factors (TFs) that regulate genetic programs (Gliwicka et al., 2013), as well as studies of callus development in other dicot crops, including cotton (Kumar et al., 2021; Wang et al., 2022; Deng et al., 2022), soybean (Zhou et al., 2023), and rapeseed, tomato, and tobacco (Iwase et al., 2013).

### RNA-Seq revealed more induced genes in OC compared to NC

In total, 3,227 DEGs were identified across the two treatments. RNA-seq analysis detected 2,086 upregulated genes in old callus (OC), compared with 1,141 in newly induced callus (NC) (Table S4; Figure 2). These results indicate broad transcriptional differences, with more genes upregulated in OC than in NC. This pattern may reflect repeated subculturing, which exposes OC to recurring stress. Repeated wounding can trigger hormonal and transcriptional reprogramming during callus formation, as reported in *Arabidopsis*. Mutant analyses also implicated cytokinin biosynthesis and an AP2 transcription factor–mediated developmental pathway in this process (Ikeuchi et al., 2017). Although OC had nearly twice as many upregulated genes as NC, many genes in both callus types encoded similar domains and may support basic callus maintenance. We also identified differentially expressed transcription factors from several families. In cotton, Li et al. (2019) examined DNA methylation across non-embryogenic and embryogenic calluses, somatic embryos, and leaves from regenerated plants. They reported methylation changes across these stages and proposed that hypomethylation associated with successive regeneration acclimation may improve regeneration capacity and contribute to cultivar improvement (Li et al., 2018). Comparable callus mechanisms have not yet been established in peanut. The gene categories identified in this study (Tables 2 and S4; Figure 3), and their implications for callus induction and maintenance, are discussed in the following sections.

### A network of genes operates in embryogenic callus growth and maintenance

The DEGs were classified into 13 categories based on available annotations. The distribution of genes across these categories suggests that callus induction in peanut involves a complex molecular network. Excluding the miscellaneous category, transcription factors (TFs) were the most abundant, while photosynthesis-related genes were the least abundant. Gene Ontology (GO) and enrichment analyses also highlighted genes involved in stress responses, carbohydrate metabolism, cell-wall metabolism, and photosynthesis (Figures 4 and 5). These processes are interconnected and likely involve the combined activity of multiple gene classes. We discuss selected classes in greater detail because of their established importance in the molecular regulation of callus development and proliferation.

#### Transcription factors and developmental regulators

The high abundance of differentially expressed TFs is expected because TFs have many functions, including roles in callus induction and organogenesis, such as shoot and root induction. Furthermore, TFs may bind to many genes involved in disease resistance, metabolism, and other processes, as they are widely known to regulate the expression of genes involved in downstream processes. One of the most important and widely recognized roles of TFs is as developmental regulators.

In our study, we identified TFs representing known DRs, including AP2/ERF (Iwase et al., 2011), WOX and LBD (Liu et al., 2018), GRAS (Kim et al., 2018), and NAC (Taoka et al., 2004), as well as TFs from families not previously identified as developmental regulators, including bHLH, WRKY, and TCP. These findings suggest that the latter may also function as developmental regulators in peanut. For instance, AP2/ERF TFs have been shown to play a major role in callus induction in *Arabidopsis* (Iwase et al., 2011, 2017). Likewise, other known DRs, such as LBD, NAC, and WOX, have diverse organogenesis-related functions, including shoot and root meristem induction and/or proliferation. These functions have been reported mostly in *Arabidopsis* (Fan et al., 2012; Hibara et al., 2003; Liu et al., 2018) and, less frequently, in other crops such as rice (Inukai et al., 2005; Zhao et al., 2009).

In peanut, a few studies have identified DRs involved in seed development (*Lec1*; Tang et al., 2018, 2024), embryo and cotyledon development (*Lec1*; Tang et al., 2024), adventitious shoot regeneration (*AhWOX13* and *AhWOX20*; Gu et al., 2026), and leaf development (*AHL23*; Liu et al., 2021). However, except for *AtLEC1* (Tang et al., 2018), the remaining genes were validated in *Arabidopsis*, despite having been identified in peanut, underscoring the need for their detailed characterization in peanut. Thus, although DRs are known, they have rarely been investigated in peanut. The putative DRs identified in the present study add to the repertoire of DRs and are potential candidates for further validation.

TFs uniquely expressed in new callus—encoding bHLH, MYB, and C2H2 zinc-finger TFs, as well as ethylene-sensitive-like proteins—may be particularly important. It would therefore be interesting to investigate whether they function as DRs and/or regulate downstream processes, such as metabolism and stress resistance, during callus induction and proliferation. TFs such as bHLH, MYB, WRKY, and NAC have also been shown to exhibit high expression during picloram-induced somatic embryogenesis in *Lilium* species (Chen et al., 2018). We also used 3 mL of 1 mg/mL picloram for callus induction; therefore, a similar scenario may have occurred in our study, resulting in high expression of these genes.

#### Auxin response genes dominate in the hormone response category

Genes in the other categories identified in this study have been widely studied in other crops, though not in peanut, for their roles in callus induction and proliferation. DEGs in the growth hormone response category mostly include genes involved in auxin response, likely reflecting the use of Picloram in our callus induction medium; Picloram is a synthetic auxin. Thus, the high expression of genes such as *GH3*, *SAUR*, and genes containing a PB1-like domain may be a response to Picloram. Supporting this interpretation, an earlier study in *Brassica napus* reported high expression of a GH3-encoding gene in response to Picloram (Ludwig-Muller et al., 2021).

#### Genes involved in oxidative stress response, methylation, kinases and seed storage

Genes involved in oxidative stress responses, methylation, and seed storage were highly upregulated in both old and new callus, indicating that they may play important roles in callus induction and development regardless of callus developmental stage. A similar study in cotton, which compared embryogenic and non-embryogenic calli, also found high expression of genes involved in oxidative stress responses and of some transcription factors (TFs) in embryogenic callus relative to non-embryogenic callus. However, methylation-related genes were downregulated in embryogenic callus in that study (Wen et al., 2020).

In the present study, as is generally the case, callus was induced from explants such as cotyledons or seeds; therefore, wounding stress was unavoidable. Oxidative stress response genes are generally expressed downstream of various stresses, including abiotic and biotic stresses. Therefore, the high expression of genes involved in oxidative stress responses, as well as disease resistance genes (e.g., those encoding NB-ARC-containing resistance proteins, TIR-domain-containing proteins, cysteine-containing leucine-rich repeats, and Gnk2-type homologs), is expected in both new and old callus. Similarly, in soybean, another oilseed crop, some genes involved in disease and oxidative stress responses were highly induced during callus induction and development (Park et al., 2023), supporting our observations.

Protein kinases are a major class of genes essential at nearly all stages of plant development, including callus formation and development. Serine/threonine-like kinases are similar to receptor-like kinases, which are activated during pathogen recognition in plants. Wound-induced callus formation also leads to downstream kinase activation. This pattern was observed in the present study, where many kinases—including serine/threonine kinases, lectin-binding receptor kinases, and wall-associated kinases (WAKs)—were highly induced. These kinases may, in turn, phosphorylate downstream targets such as TFs, metabolism-related genes, and transporters, thereby activating them. In rice, *OsSERK*, a type of receptor kinase, has been shown to regulate somatic embryogenesis and shoot regeneration in callus. Suppression of this gene by RNA interference reduced shoot regeneration, indicating its role in organ regeneration (Hu et al., 2005). In fact, kinases such as WAKs, which were uniquely expressed in new callus in the present study, are known to play an important role in cell expansion during callus formation and throughout other stages of plant development, as shown in *Arabidopsis* (Wagner and Kohorn, 2001).

#### Highly expressed genes as potentially novel DRs

To identify novel DRs in peanut, we selected the 40 most highly induced genes, with 20 from each callus type. These genes belong to different categories (Tables 2 and S4; Fig. 3) and are not necessarily TFs, which are generally known to function as DRs, as discussed earlier. DR functions may also vary depending on gene category. Examples from different plant species show that genes other than TFs can regulate callus or organ development (Table 4). However, the highly induced genes identified in both callus types require further validation to establish whether they function as novel DRs in peanut (Jiang et al., 2024; Yang et al., 2020).

We compared the chromosomal locations of these highly expressed candidate DRs with syntenic regions in soybean (Shirasawa et al., 2013) that contain QTLs associated with callus induction and embryogenesis (Song et al., 2010; Yamg et al., 2011). Across these studies, QTLs for callus induction were identified on soybean chromosomes 14 and 17, which are syntenic to peanut chromosomes A1 and B1. QTLs for somatic embryogenesis were identified on soybean chromosome 6, syntenic to peanut chromosomes A3, B3, A7, B7, A8, and B8, and on chromosome 18, syntenic to peanut chromosomes A3, B3, A4, and B4. Nearly all of the peanut chromosomes syntenic to these soybean chromosomes contain highly expressed candidate DRs that rank among the top 20 genes. Further analysis could determine whether these candidate DRs fall within peanut regions corresponding to the reported soybean QTLs—that is, whether they represent orthoQTLs.

We also compared the chromosomal distribution of DEGs with the distribution of genes on each chromosome, shown in the outer ring of the circular maps (Figure S1). The locations of 27 selected, highly expressed DEGs are marked: eight were highly expressed in NC and 19 in OC. These genes are not located in gene-dense regions, which often overlap with open chromatin regions. In addition, gene distributions on homoeologous chromosomes tend to follow similar patterns, whereas DEG distributions do not always correspond to gene density (Figure S1). This suggests that the observed DEG distribution may not be explained solely by structural features of the peanut genome and could have biological significance. The potential of these candidate morphogenic genes to improve regeneration efficiency in transformed peanut tissue should be tested. Figure 7 outlines a possible strategy for validating these candidate DRs.

**Figure 7.**
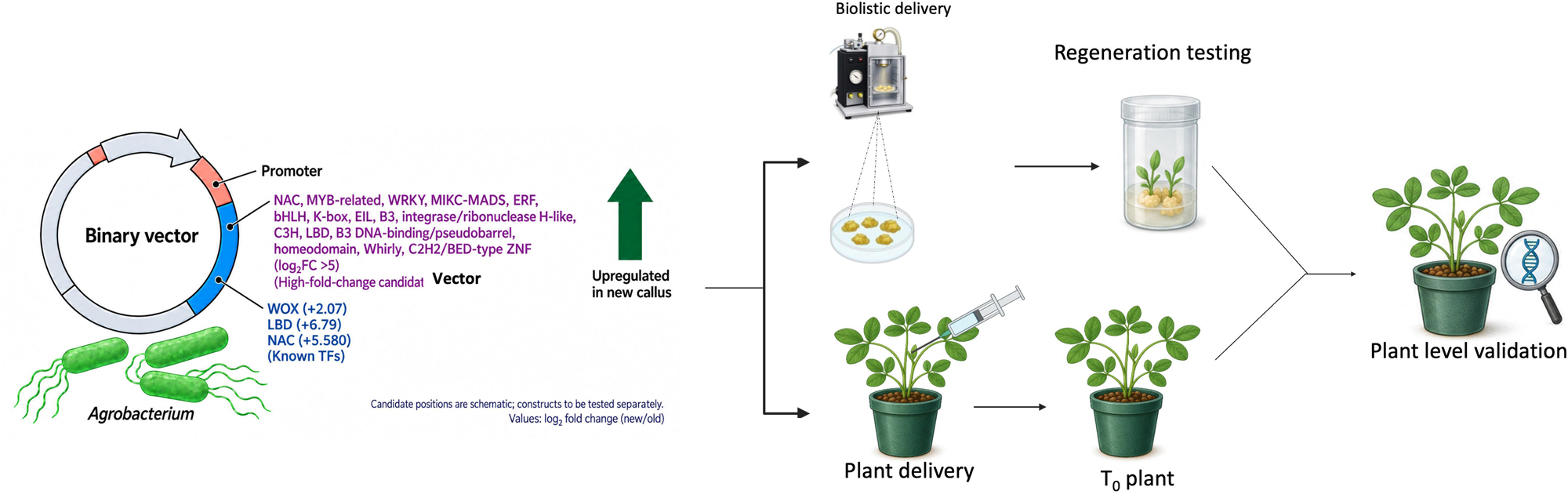
A possible workflow for validating the known and novel DRs identified in this study

## Conclusion

In this study, RNA sequencing was used to compare long-term-maintained and newly induced peanut callus tissues, providing preliminary insight into genes associated with embryogenic callus age. This work represents a first step toward understanding the molecular mechanism underlying callus development in peanut. We identified DEGs across 13 categories, including 2,086 genes highly expressed in OC and 1,141 in NC. TFs were the largest category of highly induced genes. Some have known roles in developmental regulation or morphogenesis in crops such as soybean, wheat, and rice, while other candidate DRs have not previously been associated with these processes. Overall, the findings identify candidate genes linked to callus development and provide a foundation for future studies of the molecular mechanisms of callus development and morphogenesis in peanut. The candidate DRs identified here offer promising targets for further investigation.

## Declarations

### Data availability

The raw RNA-seq data from six peanut callus samples, comprising three biological replicates each of New callus (NC) and Old callus (OC), are associated with NCBI BioProject accession number PRJNA1535505 (http://www.ncbi.nlm.nih.gov/bioproject/1535505). All other data supporting the findings of this study are included in the main text and/or Supplementary Information.

### Funding statement

This work was supported by the SC Peanut Board, the National Peanut Board, and the Foundation for Food and Agriculture Research (FFAR).

### Competing Interest Statement

The authors have declared no competing interests.

### Author Contribution Statement

**HI:** Formal Analysis, Experimentation, Investigation, Methodology, Validation, Writing-Original draft. **GS:** Writing-original draft, Writing-review & editing, Data curation and analysis. **SR**: Conceptualization, Writing-review & editing, Supervision, Funding acquisition, Resources.

## Acknowledgements

The authors would like to acknowledge SC Peanut Board, the National Peanut Board, and the Foundation for Food and Agriculture Research (FFAR) for their funding support.

## Notes

http://www.ncbi.nlm.nih.gov/bioproject/1535505

