## Supplementary Figure S1 for "Transcriptional differences between repeatedly recultured and freshly induced embryogenic calli reveal candidate developmental regulators"

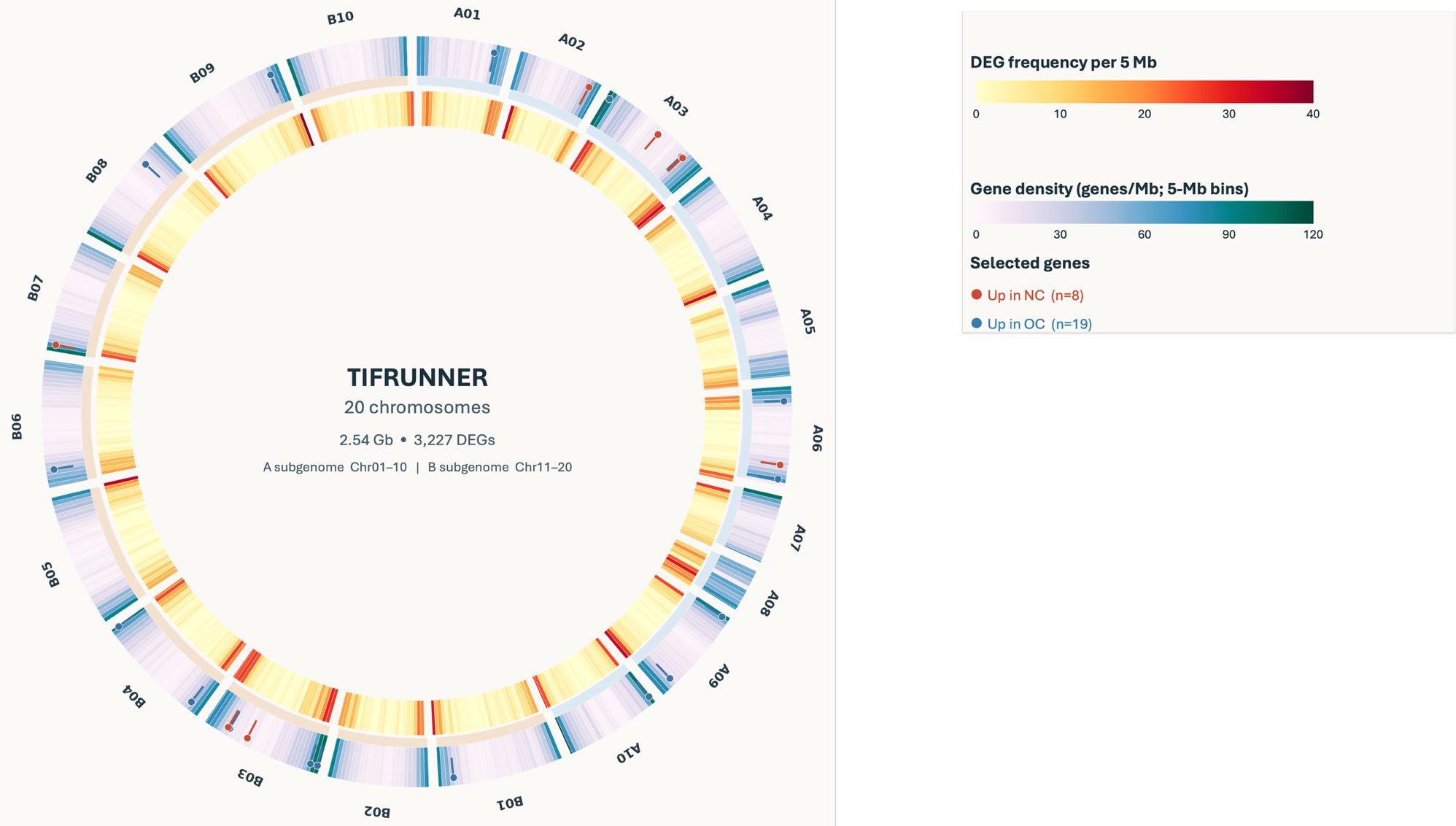

**Figure S1 Genome-wide distribution of differentially expressed genes (DEGs) and selected candidate genes in peanut.** The circular plot shows the distribution of 3,227 DEGs across the 20 chromosomes of the Tifrunner reference genome, comprising the A (A01–A10) and B (B01–B10) subgenomes. The outer heatmap represents gene density (genes/Mb), and the inner heatmap represents DEG frequency, both calculated within 5-Mb genomic bins. Colour intensities indicate increasing gene density or DEG frequency according to the respective scales. Red and blue markers indicate the genomic positions of selected genes upregulated in newly induced callus (NC; n = 8) and long-term-maintained callus (OC; n = 19), respectively.
